# Hybrid xyloglucan utilisation loci are prevalent among plant-associated Bacteroidota

**DOI:** 10.1101/2024.06.03.597110

**Authors:** Hannah Martin, Lucy A. Rogers, Laila Moushtaq, Amanda A. Brindley, Polly Forbes, Amy R. Quintion, Andrew R.J. Murphy, Tim J. Daniell, Didier Ndeh, Sam Amsbury, Andrew Hitchcock, Ian D.E.A. Lidbury

## Abstract

The plant hemicellulose xyloglucan (XyG) is secreted from the roots of numerous plant species, including cereals, and contributes towards soil aggregate formation in terrestrial systems. Whether XyG represents a key nutrient for plant-associated bacteria is unclear. The phylum Bacteroidota are abundant in the plant microbiome and provide several beneficial functions for their host. However, the metabolic and genomic traits underpinning their success remain poorly understood. Here, using proteomics, bacterial genetics, and genomics, we revealed that plant-associated *Flavobacterium*, a genus within the Bacteroidota, can efficiently utilise XyG through the occurrence of a distinct and conserved gene cluster, referred to as the Xyloglucan Utilisation Loci (XyGUL). *Flavobacterium* XyGUL is a hybrid of the molecular machinery found in gut *Bacteroides* spp., *Cellvibrio japonicus*, and the plant pathogen *Xanthomonas*. Combining protein biochemistry, computational modelling and phylogenetics, we identified a mutation in the enzyme required for initiating hydrolysis of the XyG polysaccharide, an outer membrane endoxyloglucanase glycoside hydrolase family 5 subfamily 4 (GH5_4), which enhances activity towards XyG. A subclade of GH5_4 homologs carrying this mutation were the dominant form found in soil and plant metagenomes due to their occurrence in Bacteroidota and Proteobacteria. However, only in members of the Bacteroidota spp., particularly *Flavobacterium* spp. was such a remarkable degree of XyGUL conservation detected. We propose this mechanism enables plant-associated *Flavobacterium* to specialise in competitive acquisition of XyG exudates and that this hemicellulose may represent an important nutrient source, enabling them to thrive in the plant microbiome, which is typified by intense competition for low molecular weight carbon exudates.

## Introduction

Plants provide soils with the ‘fresh’ carbon (C) required to support microbial growth, generating ‘hotspots’ of activity in regions of C deposition, such as the rhizosphere (*1, 2*). Microbial processing of plant-derived C therefore represents the entry point for new matter and energy into the microbial C pump. This biological pump determines the balance of CO_2_ liberated during aerobic respiration versus that channelled into microbial anabolism and ultimately the accumulation of recalcitrant C (*3*). Overtime, this C becomes part of the stable C pool, which is approximately 3x larger than that stored in animals and plants. Each year, soil respiration releases 10-15x more C than that emitted from anthropogenic activities (*4*). Therefore, any change in the balance of production versus respiration in response to global change will have significant ramifications for the global C cycle (*3*). Plant-derived C is partitioned into two major fractions: 1) Low molecular weight (LMW) C, which can be transformed by microbial enzyme activity within hours; and 2) Complex high molecular weight (HMW) C, e.g. glycans, which can take years to be fully degraded into their monomeric subunits (*5*). HMW C is believed to escape microbial attack, initiating the formation of soil aggregates (*5, 6*) and thus directly contributing to soil C accumulation. In addition to biogeochemical cycling, nutrient inputs have a significant influence on plant microbiome assemblage and community structure (*7*), evidenced through the impact of crop domestication (*8*).

Plant glycans (polysaccharides) are major components of plant biomass, of which hemicelluloses, such as xyloglucan (XyG), typically constitute 5-50% (*9*). Recent data has revealed XyG is a major component of root mucilage exudate. XyG is secreted at the root tip and along the entire root axes and functions to help produce the rhizosheath, a region made up largely of glycans that serves to protect roots from abrasion and desiccation (*6, 10, 11*). Through this process XyG also influences the degree of microaggregate formation, a prerequisite for soil C accumulation (*6*). Hence, these secreted HMW C polymers play an integral role in soil C storage and are likely influenced by the degree of microbial degradation. XyG binds border cells at the tip of growing roots and is an abundant component of mucilage (*12*). XyG also plays a role in regulating the severity of oomycete pathogen attack in soybean (*10*).

Historically, mycorrhizal and saprophytic fungi were considered the major plant glycan degraders, however, soil bacteria are emerging as integral players in their breakdown (*13*). In forest soils, leaf litter microbial communities are enriched with members of the phyla Pseudomonadota, Actinomycetota, and Bacteroidota (*14*). Likewise, in agricultural soil Bacteroidota and Pseudomonadota were reported as the primary consumers of cellulose, crude plant root or leaf material (*15*). Plant pathogens, such as *Xanthomonas* spp. also utilise XyG and this metabolism is considered a virulence factor, enabling the bacterium to enter plant cells (*16*). However, an understudied plant-microbe interaction is the effect of HMW C exudation on plant microbiome assemblage. This knowledge gap is driven largely by the dearth of experimentally validated genes and pathways required for hemicellulose degradation in soil bacteria, except for *Cellvibrio japonicus* (*17, 18*) and *Chitinophaga pinensis* (*19–21*).

Glycan degradation requires the possession of specialised gene sets encoding carbohydrate-active enzymes (CAZymes) to initiate degradation (*5, 13, 22*). CAZymes are categorised into broad functional groups, i.e., glycosyl hydrolases (GH) and carbohydrate esterase (CE) and are incredibly diverse (∼200 GH families), reflecting the enormous variety of naturally occurring carbohydrate structures, particularly glycans. In Bacteroidota, these gene sets are typically colocalised into discreet operons referred to as Polysaccharide Utilisation Loci (PUL) and their bioinformatic prediction has rapidly outpaced experimental validation of their precise function (*22, 23*). PUL are a hallmark of the Bacteroidota, a deep branching group of Gram-negative bacteria that specialise in HMW polymer degradation in marine and gut microbiomes (*22, 24*). Through the efficient capture of glycans, PUL provide a competitive advantage for Bacteroidota in glycan-rich environments, such as the human gut or leaf litter (*22*). Unlike their gut and marine relatives, the contribution of soil Bacteroidota towards plant or microbial glycan degradation, particularly hemicelluloses, is limited (*5, 13, 20*). Whilst *C. pinensis* can utilise a variety of glycans including several hemicelluloses, this bacterium surprisingly lacks the ability to efficiently utilise XyG (*19–21*).

*Flavobacterium*, a genus within the phylum Bacteroidota, are enriched in numerous wild and domesticated plant microbiomes relative to the surrounding bulk soil (*25–29*). Recent evidence suggests that they are one of the most metabolically active taxa in the plant microbiome, accounting for 27% of RNA reads when comprising only 6% of DNA reads (*30*). Bacteroidota are considered indicators of good soil health (*25*) and have ecological roles in suppressing various fungal and bacterial plant pathogens (*30–34*). However, their general ecological role and function remains poorly characterised in plant microbiomes, relative to other environments (*35*). Recently, we discovered *Flavobacterium* spp. have adapted to life in the plant microbiome by specialising in organophosphorus utilisation and likely play a key role in increasing phosphate availability for plants (*36, 37*). Analysing our same proteomics dataset, we further identified several CAZymes that are candidates for plant glycan utilisation, suggesting that HMW C utilisation represents a key lifestyle strategy for these bacteria (*38*).

In this study, we demonstrate *Flavobacterium* spp. are efficient utilisers of the plant hemicellulose XyG through possession of hybrid **X**yG **u**tilisation **l**oci (XyGUL). These gene clusters contained elements of the archetypal PUL identified in *Bacteroides ovatus* as well as gene clusters found in *C. japonicus* and *Xanthomonas* spp. Furthermore, we identified a XyG-specific endoglucanase associated with the XyGUL related to the glycosyl hydrolase family 5 subfamily 4 (GH5_4), subclade 2D (*39*). XyG-specific GH5_4 homologs within this clade carry a key mutation increasing their specificity and activity towards XyG. We further investigated the presence of GH5_4 homologs in soil and plant metagenomes, revealing this XyG-specialised form is prevalent in the terrestrial environment, especially in plant-associated Bacteroidota.

## Results

### Plant-associated *Flavobacterium* spp. possess a hybrid XyGUL

*F. johnsoniae* was previously reported to grow on hemicelluloses, including XyG (*40*). Therefore, we screened *Flavobacterium* sp. F52, two *Flavobacterium* spp. (OSR005 and OSR003) isolated from the rhizosphere of oilseed rape (*37*), and *F. johnsoniae* for their ability to grow on XyG, xylo-oligosaccharides (XyGO), and carob galactomannan (GalM). All four strains grew on a glucose control and on GalM and XyGOs (Figure 1a). Unlike the other three strains, *Flavobacterium* sp. F52 failed to grow on XyG, despite growing on XyGO.

**Figure 1.**
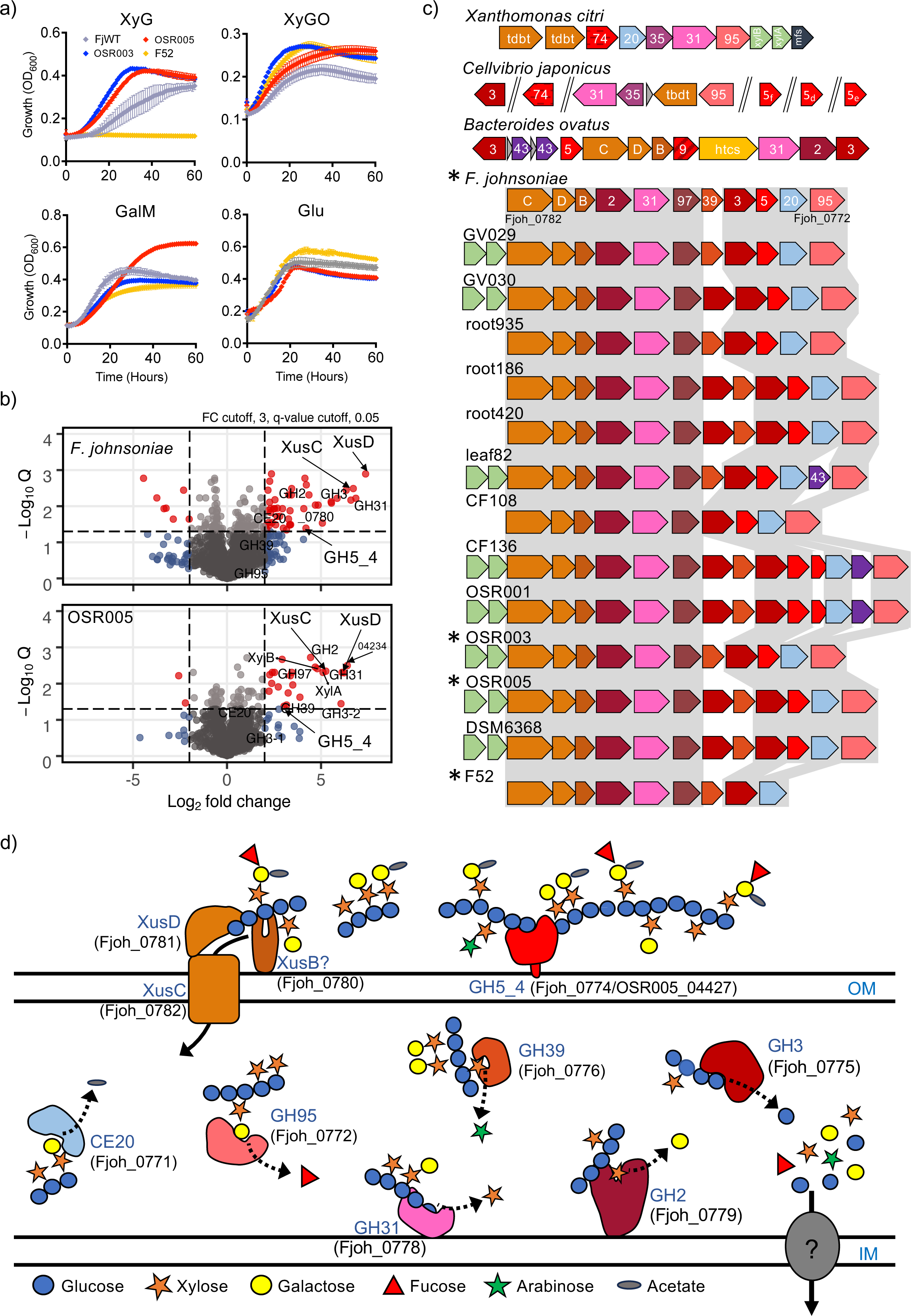
Xyloglucan utilisation by soil Bacteroidota. (a) *F. johnsoniae* grown on. plant-associated *Flavobacterium* isolated from various crop rhizospheres were grown on either glucose (Glu), galactomannan (GalM), xyloglucan (XyG) hemicelluloses, or xylooligos (XyGOs) as the sole C and energy source. Data represents the mean of triplicate cultures and error bars denote standard deviation. **(b)** Proteins enriched in the whole-cell proteomes (n=3) of either *F. johnsoniae* or *Flavobacterium* sp. OSR005 when grown on XyG compared to growth on glucose. Red data points denote statistically significant (FDR-corrected p < 0.05) proteins with greater than 2-fold enrichment. Proteins in the predicted XyGUL are highlighted. **(c)** XyGUL shares modules from *X. citri* and *B. ovatus* and are highly conserved among plant-associated *Flavobacterium* spp. (strain identifier labelled). **(d)** The predicted function and localisation of proteins encoded in the induced XyGUL with locus tags for *F. johnsoniae* provided. Colours in c and d represent the corresponding open reading frames and proteins. Locus tags correspond to *F. johnsoniae.* Numbers in d correspond to the predicted glycoside hydrolase family in the CAZY database. Asterisks represent the strain used in 1a. Abbreviations: OM, outer membrane; IM, inner membrane.

To determine the genes and proteins responsible for XyG utilisation, *F. johnsoniae* and *Flavobacterium* sp. OSR005 (*37*) were grown on either XyG or glucose and whole-cell proteomics was performed. Growth on XyG led to the significant (FDR corrected P<0.05, log_2_ FC > 2) enrichment of 68 and 74 proteins in the *F. johnsoniae* and OSR005 proteomes, respectively (Tables S4 and S5, Figure 1b). Among the most differentially synthesised proteins was a distinct PUL, hereafter named XyGUL. In *F. johnsoniae* and OSR005, XyGUL are encoded by Fjoh_0772-0782 and OSR005_04225-04238, respectively, and both contain a predicted endoxyloglucanase related to the GH5_4 family (Fjoh_0774; OSR005_4227). Surprisingly, Fjoh_0774 and OSR005_4227 showed greater sequence homology to the *C. japonicus* GH5_4 (*18*) (*Cj*GH5D, encoded by *cel5D*) than to the *Bacteroides ovatus* GH5_4 (*41*) (*Bo*GH5A, encoded by BACOVA_02653) (Table 1, Figure S1). *Cj*GH5D and *Bo*GH5A function as outer membrane-anchored XyG-specific 1-4-β-endoglucanases and are required to initiate degradation of the polysaccharide. The open reading frame (ORF) encoding this XyG-specific 1-4-β-endoglucanase was missing from *Flavobacterium* sp. F52, which harboured a truncated XyGUL (Figure 1c).

**Table 1.** Glycoside hydrolase (GH5_4) homologs used in this study, including those subjected to site directed mutagenesis.

| Enzyme name | Host strain | Locus tag | Residue (position 252) | Clade | Subclade |
| --- | --- | --- | --- | --- | --- |
| FjGH5 | *F. johnsoniae* | Fjoh_0774 | W (Trp) | 2D | I |
| 005GH5-1 | *Flavobacterium* sp. OSR005 | OSR005_04227 | W | 2D | I |
| 005GH5-2 | *Flavobacterium* sp. OSR005 | OSR005_03871 | G (Gly) | 2D | II |
| BoGH5A | *Bacteroides ovatus* | BACOVA_02653 | W | 2D | III |
| BoW252A | *Bacteroides ovatus* | BACOVA_02653 | A (Ala) | 2D | III |
| BoW252G | *Bacteroides ovatus* | BACOVA_02653 | G | 2D | III |
| CjGH5 | *Cellvibrio japonicus* | CJA_3010 | W | 2D | I |
| PaeGH5 | *Paenibacillus* sp. Root144 | 2644426200* | H (His) | 1 | NA |

\*denotes the IMG gene accession deposited in the IMG/JGI database.

Interestingly, the candidate *Flavobacterium* XyGUL combined elements of previously characterised XyGUL and associated XyG-utilising genes *in B. ovatus* (*41*)*, Xanthomonas* spp. (*16*) and *C. japonicus* (*41*). Whilst the *F. johnsoniae* GH5_4 endoxyloglucanse (*Fj*GH5) is closely related to *C. japonicus Cj*GH5D, the gene encoding *Cj*GH5D is located away from the truncated and fragmented gene cluster required to import and degrade XyGOs (*18, 41*) (Figure 1c). Other XyGUL-encoded proteins predicted to sequentially hydrolyse the XyG oligosaccharides were annotated as L-α fucosidase (GH95), β-glucosidase (GH3), 1,4-β-xylosidase (GH39), α-glucosidase (GH97), α-D-xyloside xylohydrolase (GH31), β-galactosidase (GH2), a SusCD-like outer membrane transport system (hereafter termed XusCD), and the distinct O-acetylesterase (CE20) that was recently characterised in *Xanothomonas* (*16*) (Figure 1c & d). XusC and D were the most differentially abundant proteins during growth on XyG (Figure 1b). The *Flavobacterium* XyGUL showed high degree of conservation and synteny across all plant-associated strains analysed (Figure 1c), in contrast to the XyGUL found in *Bacteroides.* spp. (*41*), with no rearrangements and only few instances of gene insertions. GH39, predicted to hydrolyse the Xyl(α*1-2*)Araf linkage found in *solanaceous* plants, such as tomato, was only present in *Flavobacterium*. Together, these data suggest *Flavobacterium* harbour a specialised XyGUL capable of capturing and breaking down XyG from various plant species.

### XyGUL encoded proteins are essential for efficient growth on XyG in *F. johnsoniae*

To determine the *in vivo* contribution of XyGUL encoded proteins to growth on XyG, two knockout strains of *F. johnsoniae* were generated. The first had a deletion of *fjoh_0774*, encoding the GH5 enzyme predicted to initiate depolymerisation of the XyG polysaccharide, and the second was an *fjoh_0781-2* mutant lacking the XusCD system predicted to be required for oligosaccharide uptake (Figure 1d). The isogenic wild-type parent and both mutant strains grew comparably on either glucose or GalM, but, unlike the wild type, Δ*0774* was unable to grow on XyG, whilst the growth of the Δ*0781-2* mutant was significantly curtailed (Figure 2). These data are consistent with the proteomics analysis (Figure 1b), demonstrating that the predicted XyGUL is essential for growth on XyG (Figure 2). Complementation of each mutant with an *in trans* copy of the respective gene(s) restored their ability to grow on XyG (Figure 2). As expected, the Δ*0774* mutant, lacking the outer membrane initiator enzyme *Fj*GH5, was capable of growth on commercially synthesised on XyGOs (Figure 2). However, Δ*0781-2* also grew on XyGOs, albeit at a slower rate, in contrast to its phenotype on XyG (Figure 2). This suggests that either *Fj*GH5 and XusCD interact for efficient hydrolysis of the polysaccharide backbone prior to import, or that XyGOs produced by *Fj*GH5 are not the same as those present in the hepta-, octa-, nona-saccharides commercial mix (MEGAZYME), and that other SusCD-like complexes can import the latter.

**Figure 2.**
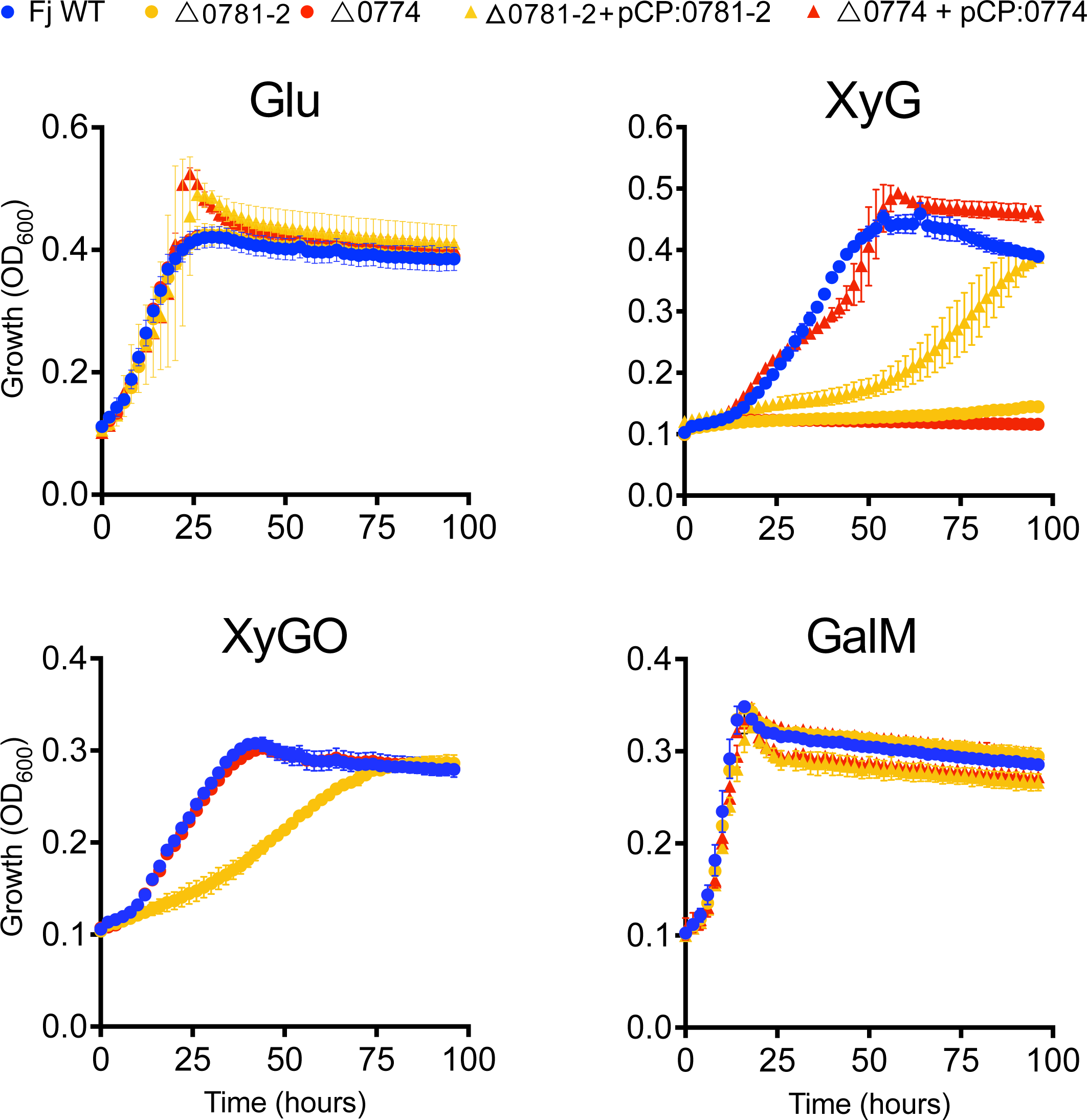
Genetic basis of xyloglucan utilisation in *Flavobacterium johnsoniae*. The wild type (blue circles), the outer membrane GH5_4 endxyloglucanse (Δ*0774*) mutant (red circles), the outer membrane TonB-dependent transporter and cognate lipoprotein (Δ*0781-2*) mutant (yellow circles) were grown on either glucose, XyG, XyGO, or GalM as the sole C and energy source. Both mutants were complemented (triangles) with their respective native genes. Growth assays were performed in triplicate and error bars denote the standard deviation from the mean.

### Microdiversity of GH5_4 homologs in *Flavobacterium* spp. suggests functional diversification

In several plant-associated *Flavobacterium* spp., BLASTP identified multiple ORFs encoding GH5_4 homologs. Phylogenetic reconstruction of these homologs alongside *Bo*GH5, *Cj*GH5d, and other previously characterised GH5_4 homologs eliciting mannanase, xylanase, and glucanases activity revealed the presence of two distinct *Flavobacterium* GH5_4 groups (Figure 3a). These two subgroups (Type I and Type II) shared greater similarity to each other than to the archetypal *Bo*GH5A. Whilst 7/8 residues, previously shown to be involved in XyG hydrolysis in *Bo*GH5A (*41*) were conserved across Type I and Type II homologs, residue Trp252 in *Bo*GH5A was not (Figure 3a & S1). Trp252 is conserved in all Type I homologs, including *Fj*GH5, the GH5_4 enzyme encoded by OSR005_04227 in *Flavobacterium* sp. OSR005 (Figure 1), hereafter termed 005GH5-1 (Table 1), and *Cj*GH5D (*17*). In the majority of Type II homologs, Trp252 is replaced with either Ala or Gly. The genes encoding type I homologs are all found in XyGUL, however the genes encoding Type II GH5_4 homologs were all found in distinct PUL (XyGUL2 in Figure 3b). This was confirmed by increasing the number of plant-associated *Flavobacterium* genomes screened, including the addition of MAG retrieved from plant rhizosphere metagenomes (Table S3). Even the genes encoding the few Type II forms carrying the Trp residue (Figure S1) were found in XyGUL2-like PUL. XyGUL2 is present in fewer *Flavobacterium* genomes and has far less gene synteny and conservation than XyGUL1 (Figure 3c). These alternative PUL contain ORFs for various exo-acting GHs, distinct SusCD-like systems, and in some cases a GH74 homologue similar to the endoxyloglucanase recently shown to be functional in *Xanthomonas* spp. (*16*). In addition, *Flavobacterium* sp. OSR005, harbours a Type II GH5_4 (hereafter referred to as 005GH5-2), encoded by OSR005_03871, which contains an Ala in place of the aforementioned Trp252 residue. Neither 005GH5-2 nor other XyGUL2 proteins were detected during growth on XyG (Figure 1b, Table S5), suggesting they do not play a role in XyG utilisation in *Flavobacterium* sp. OSR005.

**Figure 3.**
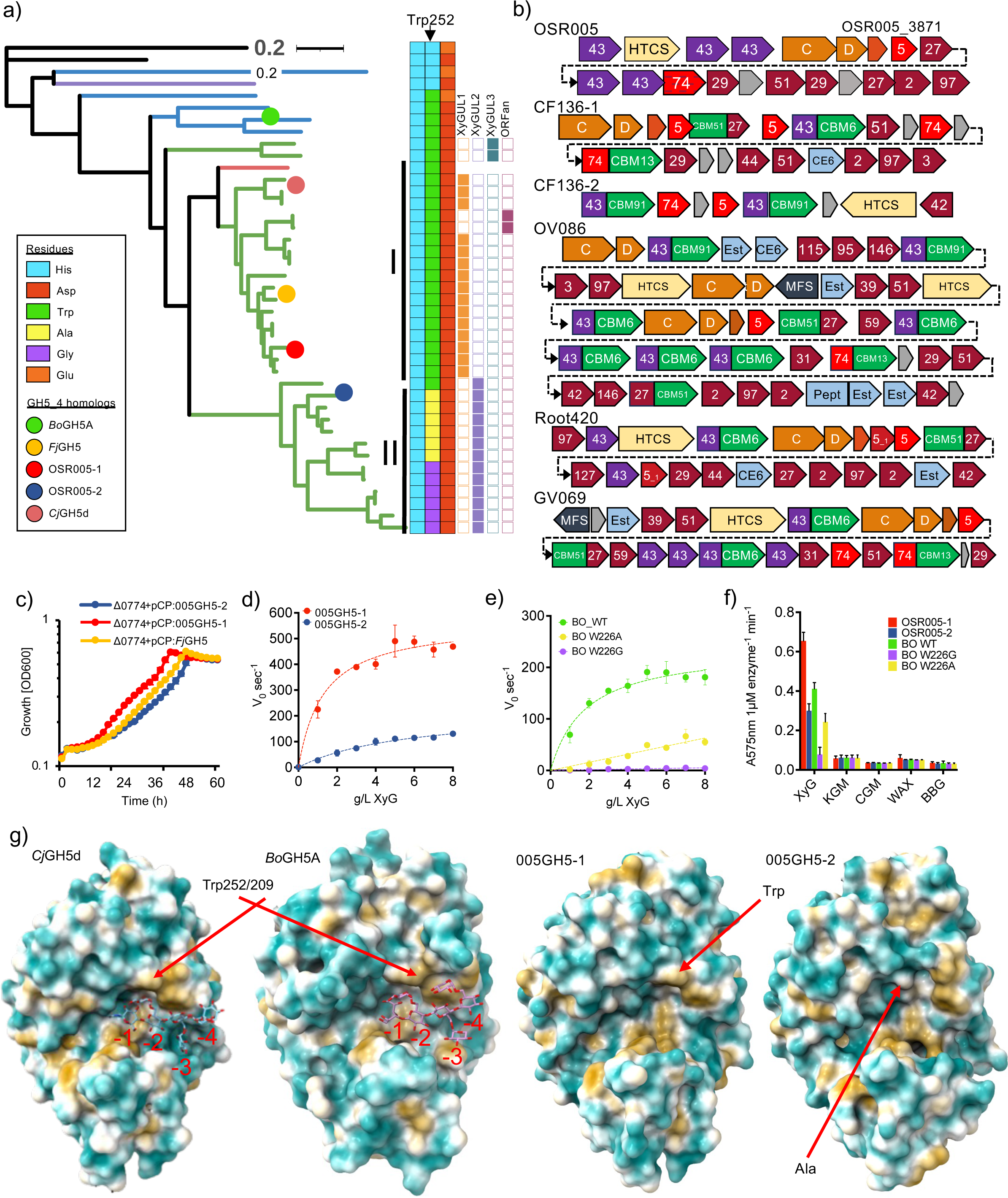
Characterisation of GH5_4 homologs in *Flavobacterium* spp. **(a)** Phylogenetic reconstruction of GH5_4 homologs identified in *Flavobacterium* spp. alongside those previously characterised, showing the variable Trp252 residue (*Bo*GH5A) and each adjacent amino acid residue. The genomic localisation of the GH5_4 homologs is given in columns to the right of the residues. Note, the Trp-containing forms in *Flavobacterium* (green branches) are almost exclusively associated with XyGUL. I and II represent the identified Type I and Type II *Flavobacterium* GH5_4 homologs. Abbreviations: Est, esterase; Pept, peptidase; CBM, carbohydrate binding module; MFS, major facilitator superfamily transporter; HTCS, hybrid two component sensor **(b)** Predicted PUL (XyGUL2) containing either Ala or Gly containing GH5_4 homologs in *Flavobacterium* spp. demonstrating distinct CAZyme organisation relative to XyGUL1. Numbers denote glycoside hydrolase family predictions. **(c)** Growth (n=3) of the *F. johnsoniae* Δ*0774* mutant complemented with either its native gene homolog or the two *Flavobacterium* sp. OSR005 GH5_4 homologs (005GH5-1, 005GH5-2). **(d-e)** Enzyme kinetics for DNSA assays were performed to determine endoxyloglucanase activity (tamarind XyG) of purified recombinant GH5_4 homologs from *Flavobacterium* sp. OSR005 **(d)** and *Bo*GH5A wild type (WT), W252A and W252G variants **(e)**. These five recombinant GH5_4 enzymes were also tested for activity against Konjac-glucomannan (KGM), Carob-galactomannan (CGM), wheat arabinoxylan (WAX), and barley beta-glucan (BBG) **(f)**. Reactions (n=3) were stopped after 10 min and reducing sugar ends were quantified (Abs575 nm) and normalised to 1 µM enzyme per min. **(g)** *Cj*GH5d (pdb: 5oyd) and BoGH5A (pdb: 3zmr) structures depicting surface hydrophobicity determined by X-Ray crystallography modelled with XyG bound visualising the stacking interaction between the third xylose side branch found in a typical XXXG motif, including those occurring with Trp252/209 at the -2 subsite. Alphafold2 generated models of 005GH5-1 and 005GH5-2. Arrows indicate the Trp residue that is substituted with Ala in 005GH5-2. Enzyme and growth assays were performed in triplicate and error bars denote the standard deviation from the mean.

To determine if Type II GH5_4 homologs were functional, we complemented the *F. johnsoniae* Δ*0774* mutant with the genes encoding 005GH5-1 and 005GH5-2 expressed from the constitutive *ompAFj* promoter. Both 005GH5-1 and 005GH5-2 restored the ability of Δ*0774* to grow on XyG as the sole C source, with the 005GH5-1 strain showing a greater initial growth rate and 005GH5-2 the slowest (Figure 3c). To test if the lower growth rate observed for 005GH5-2, which carries the W252A substitution, was due to a lower enzyme activity, we purified recombinant OSR005-1, 005GH5-2 and the archetypal *Bo*GH5A following heterologous over-production in *E. coli*. Recombinant 005GH5-1 had a significantly greater turnover rate (*K*_cat_ = 566.2 min^-1^) than recombinant 005GH5-2 (*K*_cat_ = 223.5 min^-1^) and a lower *K*_m_ (OSR005-1 = 1.3 mg ml^-1^, OSR005-2 = 5.7 mg ml^-1^) (Figure 3d). Recombinant *Bo*GH5A modified with either W252A or W252G substitutions replicated this dramatic reduction in endoxyloglucanase activity (Figure 3e), with W252G having the greatest reduction, requiring 10x more enzyme to detect observable activity (Figure S2). Neither OSR005-1, OSR005-2, *Bo*GH5A, *Bo*W252A nor *Bo*W252G conveyed substrate promiscuity towards other glycans typically found in the plant microbiome (Figure 3f). Based on structural homology modelling and previous structural data for *Bo*GH5A and *Cj*GH5d (*18, 41*), Trp252/209 interacts with the xylose residue occupying the -2 glucose position in XX**(X)**G-type saccharides, such as tamarind XyG (Figure 3g; Figure S3). This would explain why mutation of Trp252 results in the observed decrease in activity. Modelling the surface hydrophobicity revealed that possession of Trp252 likely generates a stacking interaction which may stabilise the docking of XXXG-type XyG. In 005GH5-2 and other promiscuous GH5_4 enzymes where Trp252 is absent, a clear cavity is present that would significantly reduce this stacking interaction between the aromatic residue and the xylose occupying the -2 subsite (Figure 3g; Figure S3). Taken together, these data reveal Type II GH5_4 homologs may have subsequently evolved to specialise on another glycan or variation of XyG, perhaps the XXGG-type typical of solanaceous plants.

### GH5_4 homologs are enriched in plant-associated Bacteroidota genomes

Next, we investigated if XyG utilisation in *Flavobacterium* is an adaptation to life in the plant microbiome by analysing our previous database containing ∼100 genomes representing *Flavobacterium* spp. isolated from distinct ecological niches (*37*). In addition to searching for GH5_4 homologs, we also searched for homologs related to other XyGUL components and candidate GH10 endoxylanases (pfam00331) required to hydrolysis xylan backbones (*42*). Xylan is another hemicellulose secreted from plant roots (*43*). ORFs encoding XyGUL components were more prevalent among plant-associated and closely related strains (Figure 4a). Likewise, GH10 homologs followed a similar pattern. Several plant-associated *Flavobacterium* strains sometimes possessed up to six closely related GH5_4 homologs, each associated with either Type I or II. The most prevalent were the canonical GH5-1 forms found in the XyGUL, followed by homologs related to 005GH5-2 (group GH5-3 in Figure 4a). Some *Flavobacterium* spp. possess a second Type I GH5_4 homolog (GH5-2 in Figure 4a), typically located adjacent to the GH5-1 in the XyGUL (e.g. CF136 and OSR001 in Figure 1d).

**Figure 4.**
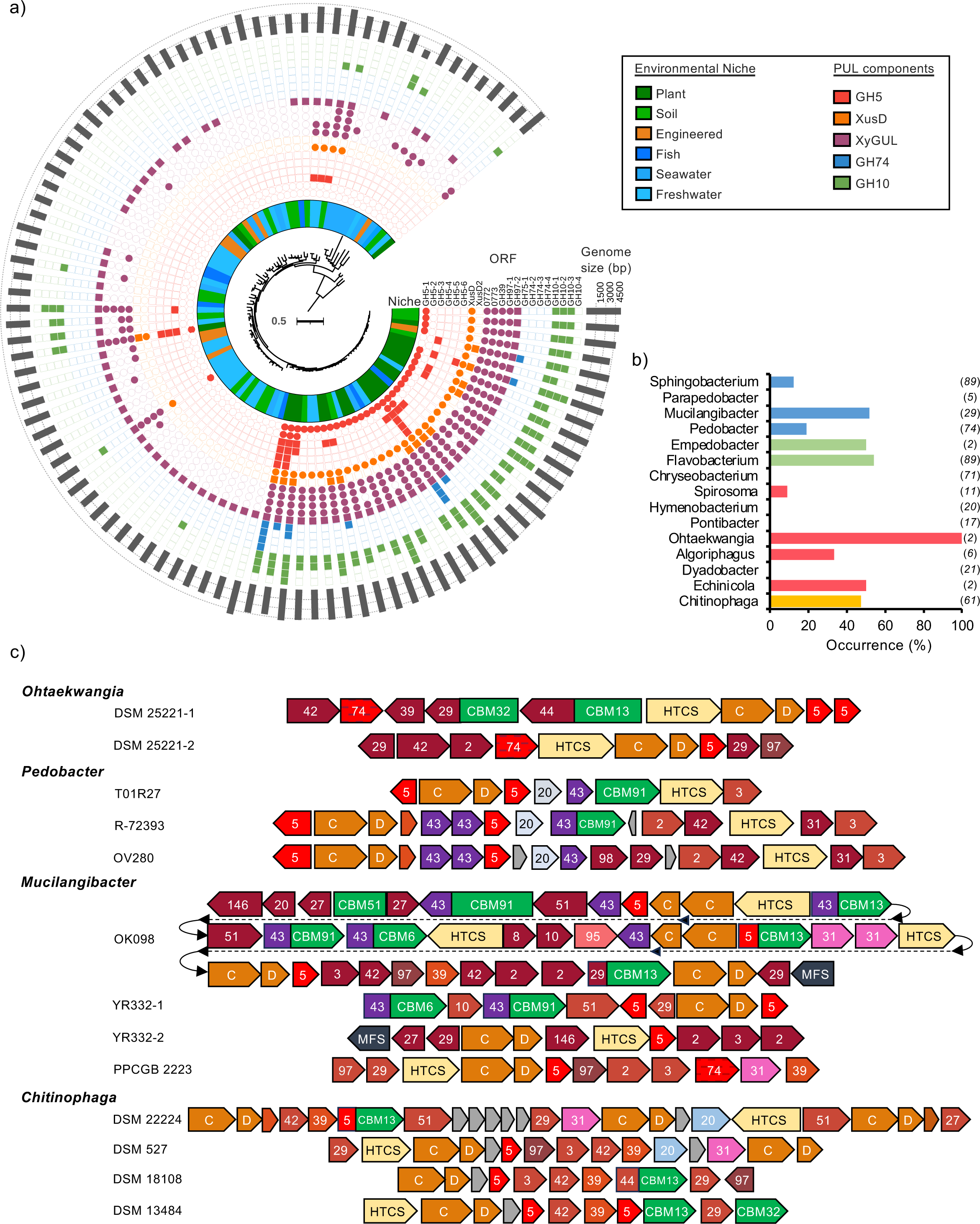
The occurrence and diversity of GH5_4 homologs in terrestrial Bacteroidota spp. **(a)** Phylogenomic analysis of our previously generated multi-loci maximum-likelihood consensus tree, inferred from the comparison of 10 housekeeping and core genes present in 102 Flavobacterium isolates (*37*). The presence (filled symbol) or absence (hollow symbol) of CAZyme ORFs associated with PUL are displayed, as well as the genome size of each isolate (outer ring). The inner ring denotes the environmental niche the genome was isolated. **(b)** The prevalence of GH5_4 homologs in the genomes from different genera within the phylum Bacteroidota, determined through BLASTP (cut off, e^-40^). The number of genomes screened per genus is given in the parentheses. Colours denote the associated class rank. **(c)** Selected PUL containing GH5_4 homologs identified in other Bacteroidota spp. Numbers denote glycoside hydrolase family predictions. Abbreviations: CBM, carbohydrate binding module, HTCS, hybrid two component sensor. Colour schemes as per previous figures.

*C. pinenesis* DSM2558 cannot efficiently grow on XyG (*19–21*) and BLASTP confirmed that this strain lacks either a GH5_4 homolog or a XyGUL. To determine whether XyGUL is restricted to plant-associated *Flavobacterium* or found within the Bacteroidota phylum more widely, we screened genomes deposited in the IMG/JGI database (Table S2) for the presence of GH5_4 homologs. Genomes were restricted to those retrieved from terrestrial environments, i.e., soil and plant, and encompassed *Chitinophagaceae*, *Sphingobacteraceae, Flavobacteriaceae*, and *Cytophagaceae*. We detected both inter- and intra-genus variation in the occurrence of GH5_4 homologs in the genomes of Bacteroidota spp. (Figure 4b). The highest percentage of genomes possessing GH5_4 homologs belonged to *Flavobacterium* (54%), with almost all plant-associated strains possessing the gene cluster. Despite belonging to the family *Flavobacteriaceae*, we found no GH5_4 homologs in plant-associated *Chryseobacterium*. Likewise, no GH5_4 homologs were found in the *Pontibacter* and *Hymenobacterium* genomes we screened. Genomes related to both *Mucilaginibacter* (51%) and *Chitinophaga* (46%) also had a relatively high number with at least one GH5_5 homolog present.

Given that several genomes possessed multiple GH5_4 homologs, we performed phylogenomics to determine whether they belonged to Type I or Type II forms (Figure S4). Most GH5_4 homologs identified in non-*Flavobacterium* Bacteroidota fell into the Type II subgroup. However almost all harboured the Trp residue, except for a few containing Tyr, and some being closely related to the Ala- and Gly-harbouring Type II forms. Genomes from the class Sphingobacteriia often possessed two or more homologs. Two major clusters of *Chitinophaga* were present, all harbouring the Trp residue, and these were typically mutually exclusive within genomes and found in distinct PUL. Interestingly, no Bacteroidota genomes possessed only a Type II GH5_4 carrying the Ala or Gly mutation, strengthening the hypothesis that this form has an auxiliary role in XyG hydrolysis. Taken together, whilst there is a large diversity of GH5_4 and XyGUL-like clusters, whether these are all functional as part of dedicated XyG utilisation pathways remains uncertain.

In other Bacteroidota spp. the organisation of PUL harbouring Type II GH5_4 homologs carrying the Trp residue differed substantially from the *Flavobacterium* XyGUL (Figure 4c). These PUL resembled the organisation and features, such as carbohydrate binding domains (CBMs), associated with the XyGUL2 cluster found in *Flavobacterium* spp., which was not induced during growth on XyG in OSR005 (Figure 1b, Table S5). Therefore, whether the XyGUL-like clusters identified in other Bacteroidota genera also specialise in XyG utilisation remains an open question.

### GH5_4 subclade 2D has radiated in soil and plant microbiomes

The GH5_4 family has recently been structured into three main clades (named I, II, III) and subclades (*44*), with *Bo*GH5A and *Cj*GH5d belonging to subclade 2D (*44*). Given the high prevalence of GH5_4 homologs in plant-associated Bacteroidota, we performed BLASTP on over 700 plant/soil metagenomes deposited in the IMG/JGI database (Table S3). Two GH5 sequences were used as queries: *Fj*GH5 (Fjoh_0774) and a GH5_4 from *Paenibacillus* sp. Root144 (IMG gene id, 2644426200), the latter closely related to a commercial *Paenibacillus* endoxyloglucanase (Megazyme) and represents GH5_4 homolog from subclade 1. All environmental ORFs retrieved (n=7636) were locally aligned (BLASTP) to all GH5 enzymes in the CAZYdb (n = 1123) (*45*). In total, 7136 ORFs aligned to 254 ORFs from CAZYdb and were all related to the GH5_4 subfamily. Homologs related to Bacteroidota (N= 39150) and Proteobacteria (n= 39031) constituted much of the diversity found in soil (Figure S5). At the genus-level, homologs related to *Capsulimonas* (Actinomycetota, n=11783) and *Flavobacterium* (n=11232) were the most abundant, followed by *Cellvibrio* (8700), *Mucilaginibacter* (n=8697), and members of the family *Chitinophagaceae* (*Pseudobacter*; n =7967, *Chitinophaga*; n=5516).

Phylogenetic reconstruction revealed most environmental homologs were related to GH5_4 clade 2, with most sequences belonging to subclade 2D. This subgroup contains *Fj*GH5, 005GH5-1, *Cj*GH5d, and all homologs related to Bacteroidota, including *Flavobacterium* (Figure 5b). Meanwhile, GH5_4 homologs related to Gram-positive bacteria, primarily, Actinomycetota and Bacillota, were found in subclades 1 and 2. As observed for *Flavobacterium* Type I and Type II GH5_4 homologs, the eight residues involved in XyG binding and hydrolysis by *Bo*GH5A (*41*) were highly conserved between clades I, II, and III, again with the exception of Trp252. This residue was predominantly substituted with either His or Gly in clades 1, 3, and subclades 2A, 2B, and 2C (Figure 5b). Despite the occurrence of W252A- and W252G-GH5_4 Type II forms in isolates related to several Bacteroidota genera, in soil/plant metagenomes only *Flavobacterium* Type II forms were detected. Most GH5_4 homologs related to other Bacteroidota and Proteobacteria spp. were Type I. Together, these data demonstrate subclade 2D has radiated in soil and become the dominant form. Furthermore, these analyses highlight a possible role for horizontal gene transfer of the GH5_4 enzyme between Bacteroidota and Proteobacteria, such as *Cellvibrio*, in response to occupying a similar niche.

**Figure 5.**
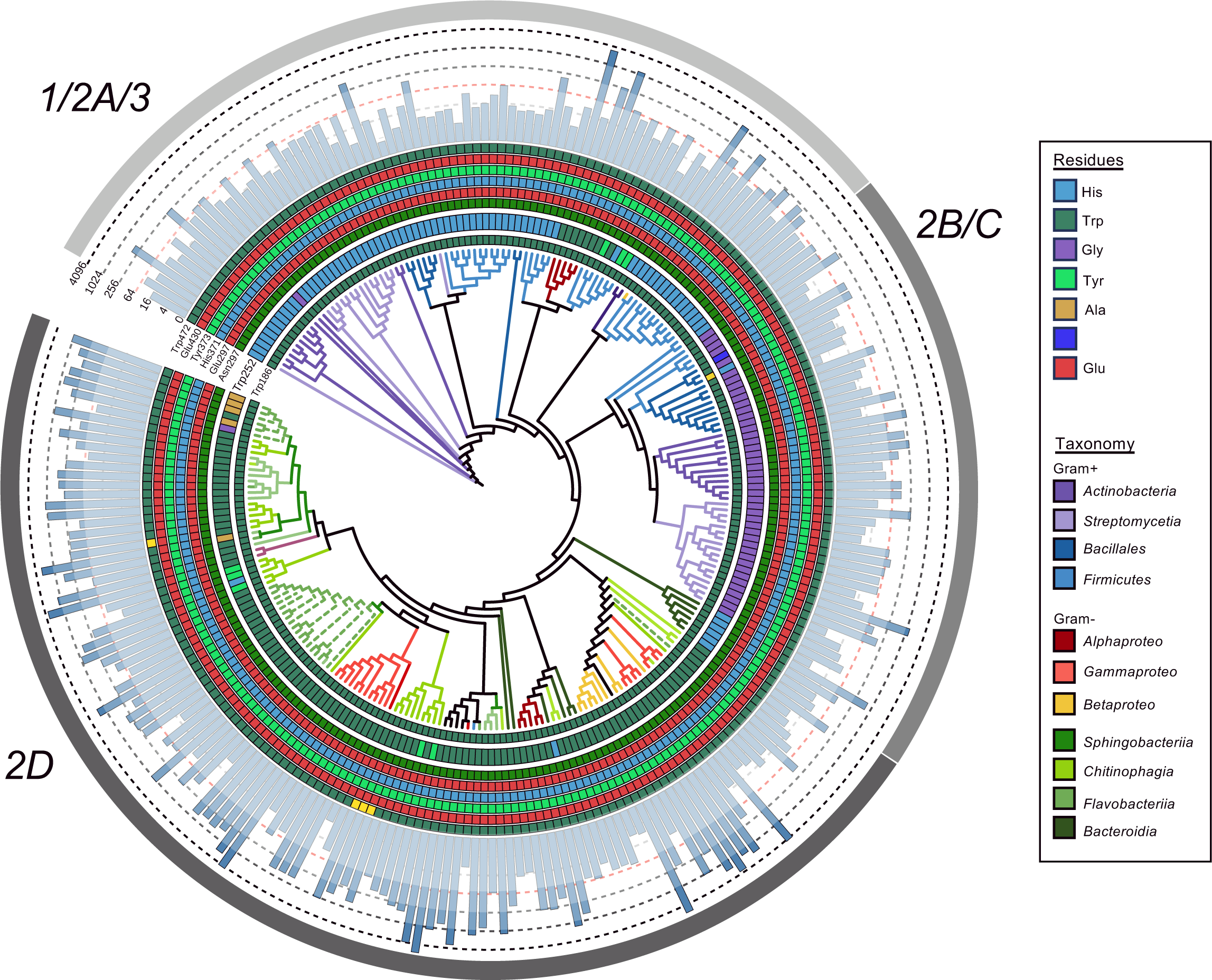
Distribution of GH5_4 homologs in soil- and plant-associated metagenomes. Reconstructed phylogeny (maximum likelihood method, bootstrap 1000) of GH5_4 homologs in the CAZyme database that best represent the ORFs retrieved from the metagenomes. The amino acid present at each of the key residue sites experimentally determined in previous studies are presented as coloured rings. The outer bar plots represent the overall gene abundance across all metagenomes. Branches are coloured based on their taxonomic classification at the class level. The outer ring represent the GH5_4 clades (I,II, or III) previously identified by (*44*).

## Discussion

Here, we demonstrate *Flavobacterium* spp. can efficiently utilise the plant hemicellulose XyG and, through the identification of molecular markers, show that this metabolic trait is prevalent in plant-associated Bacteroidota spp. The explosion of next-generation sequencing studies investigating the composition of plant microbiota has revealed Bacterdoidota, particularly *Flavobacterium*, are highly enriched in this niche (*27, 28, 46, 47*). However, these bacteria are typically not enriched when LMW substrates, such as sugars, organic acids, aromatics and phenolics, are supplemented to soil samples under laboratory conditions (*48–50*). The possession of XyGUL and similar hemicellulose utilisation systems may therefore provide Bacteroidota with a competitive advantage when invading and persisting in the plant microbiome, facilitated through resource diversification (*24*). The high prevalence of XyGUL in plant-associated genomes and low prevalence in those retrieved from other environmental niches, such as seawater, further suggests a strong selection for XyG utilisation as a strategy to succeed in the plant microbiome, similar to their organophosphorus utilisation capabilities (*37*). These data are also consistent with previous comparative genomics analyses that indicated terrestrial *Flavobacterium* have a greater ratio of GH enzymes relative to peptidases, including those predicted to target plant pectins (*51*). Whilst Gammaproteobacteria, such as *Cellvibrio* and *Xanthomonas*, possess endo-acting xyloglucanases, these bacteria only contain a TonB-dependent transporter, akin to XusC (*16, 17*), lacking the surface-exposed glycan binding domain (XusD) identified in this study. Therefore, possession of XusCD may increase the competitive ability of *Flavobacterium* to capture these complex exudates (*52*), consistent with the ecological function of these transporters in marine and gut microbiomes (*53, 54*).

Terrestrial Bacteroidota can utilise other HMW substrates, including pectin (*55*), alternative hemicelluloses (*13, 20, 21*), fungal polysaccharides (*19, 56*) and alternative plant cell wall components (*13, 21*). Together with our data, these observations support a model whereby HMW C is the preferential nutrient and energy source for Bacteroidota in soil and plant microbiomes. The domestication of agricultural crops is driving a significant loss of various key microbiota, including Bacteroidota, hypothesised to be a consequence of changes in crop root exudation profiles with a relative increase in the ratio of LMW:HMW (*8*). This reduction in beneficial microbes, such as *Flavobacteraceae* and *Chitinophagaceae*, may have negative impacts on agricultural soil health (*57*) and the plants ability to suppress pathogens (*31, 32, 58–60*). Interestingly, the relative abundance of genes encoding XyGUL components, such as GH5, GH31, GH3, and GH95 were also significantly higher in healthy versus diseased pepper plants, when challenged with *Fusarium* (*61*). Collectively, these studies and ours highlight a possible link between Bacteroidota, HMW C utilisation, and plant disease suppression. We propose, future research should focus on explicitly linking the connection between HMW exudation and the assemblage of Bacteroidota in the plant microbiome in the context of crop domestication and host disease These studies are essential to better understand the drivers of Bacteroidota assemblage and host-microbe interactions in the plant microbiome (*35*).

Given the proposed importance of plant polysaccharides in soil aggregation and the long-term storage of C (*3, 6*), degradation of these molecules may represent a significant and relatively overlooked cog in the global C cycle. The comparatively efficient utilisation of glycans by Bacteroidota relative to non-Bacteroidota, as observed in marine systems (*52, 62, 63*), may therefore have consequences for the microbial C pump (*3*), which can be altered by changes in bacterial C use efficiency (*64, 65*). Microbial polysaccharides also represent a major fraction of recalcitrant or ‘stabilised’ C in soil, a fraction which is vulnerable to microbial attack in response to a climate-induced influx of labile C or changes in land-use intensity (*3, 4, 64, 65, 66*). Whether shifts in Bacteroidota abundance and diversity, which are known to be good indicators of soil health, could influence this key step in the terrestrial global C cycle warrants further investigation (*8, 57*).

The lack of XyGUL in certain genera related to Bacteroidota, e.g., *Chryseobacterium*, coupled with a 10-60% occurrence of GH5_4 homologs in other Bacteroidota genera, suggests some level of functional partitioning within this phylum. Indeed, *Chryseobacterium* spp. possess an enhanced capability to degrade microbial polysaccharides associated with Gram-positive peptidoglycan compared to *F. johnsoniae* or *Sphingobacterium* sp. (*67*). *C. pinensis* also lacks the ability to utilise XyG despite its capability to grow on other hemicelluloses and fungal glycans (*19–21, 68*) and our comparative genomics confirmed this bacterium lacks a GH5_4 homolog. Hence, whilst Bacteroidota likely specialise in HMW C utilisation *in situ*, resource partitioning or metabolic heterogeneity within this phylum exists to target different HMW C substrates.

Our data also reveals subclade 2D of the GH5_4 has radiated in soil microbiomes and is the dominant form, in contrast with the abundant forms found in engineered systems or animal guts (*44*). Clade 2D carried a distinct mutation at Trp252 (position in *Bo*GH5A), which is typically Gly, Ala or His in clades 1 and 3. Clade 3 GH5_4 enzymes possess high activity towards multiple polysaccharides in addition to xyloglucan, in contrast to clade 2D homologs produced by *C. japonicus* (*Cj*GH5d, *Cj*GH5e, *Cj*GH5f) (*18, 39, 44*). Hence, the presence of clade 1 and 3 GH5_4 enzymes in Actinomycetota and Bacillota may reflect a trade off whereby these bacteria carry fewer CAZymes with greater individual substrate ranges relative to Bacteroidota in order to scavenge complex C molecules in bulk soil away from plant roots (*5, 24, 69, 70*). However, enzyme specificity versus promiscuity is likely driven by many more mutations that influence active site architecture through alterations in secondary structure (*39*). This may explain why mutation of Trp252 *Bo*GH5A did not broaden its substrate range. In gut Bacteroidota, distinct PUL are required to degrade simple and complex arabinoxylans, which are differentially regulated in response to these different forms of the polysaccharide (*23, 71*). The existence of Type II GH5_4 homologs carrying a single mutation and typically found in PUL that significantly differ from the conserved *Flavobacterium* XyGUL in their organisation and overall complexity may present something similar. Indeed, XyG is often part of a larger polysaccharide exudate complex, which includes pectin and xylan complexes (*11*). These complex Type II-harbouring PUL may therefore represent specialisation in utilising either non-exudate plant polysaccharides, particularly those associated with plant cell walls or root tip border cell-mucilage matrices (*9, 21*) or more complex forms released by plant roots (*11*).

In summary, using *Flavobacterium* as the model, we identified highly conserved XyGUL among plant-associated members of this genus. Whilst the initiator enzyme for XyG polysaccharide hydrolysis, GH5_4, is found in the genomes other Bacteroidota and Proteobacteria spp., we hypothesise the specialised *Flavobacterium* XyGUL, harbouring the active Type I form, enables these bacteria to competitively acquire this complex carbohydrate. Given the emergent knowledge that most plants, including globally important crop species, exude significant quantities of XyG, we propose this hemicellulose may present an important nutrient source for plant-associated *Flavobacterium* and underpins their ability to successfully invade and persist in a highly competitive plant microbiome.

## Materials and methods

### Bacterial strains and growth medium

*F. johnsoniae* UW101 (DSM2064) was purchased from the Deutsche Sammlung von Mikroorganismen und Zellkulturen (DSMZ) collection. *Flavobacterium* sp. OSR005 and *Flavobacterium* sp. OSR003 were isolated previously (*37*). *Flavobacterium* sp. F52 was kindly donated from the Cytryn lab (*71*). *Flavobacterium* strains were routinely maintained on casitone yeast extract medium (CYE) (52) containing casitone (4 g L^-1^), yeast extract (1.25 g L^-1^), MgCl_2_ (350 mg L^-1^), and agar (20 g L^-1^). For conjugation experiments, MgCl_2_ was substituted with CaCl_2_ (1.36 g L^-1^). For growth experiments investigating hemicellulose degradation, *Flavobacterium* strains were grown in a modified minimal A medium (*72, 73*), containing NaCl (200 mg L^-1^), NH_4_Cl (450 mg L^-1^), CaCl_2_ (200 mg L^-1^), KCl (300 mg L^-1^) and MgSO_4_ (350 mg L^-1^). After autoclaving, filter-sterilised yeast extract (10 mg L^-1^), FeCl_2_ (2 mg L^-1^), MnCl_2_ (2 mg L^-1^), NaH_2_PO_4_ (100mg L^-1^) and 20mM HEPES buffer pH7.4 were added. This medium was supplemented with either 0.25-0.4% (w/v) glucose, tamarind XyG (Megazyme, CAS Number: 37294-28-3), Xyloglucan oligosaccharides (hepta+octa+nona saccharides, Megazyme, CAS Number: 121591-98-8) or Carob galactomannan (Megazyme, CAS Number: 11078-30-1) as the sole C source. Growth assays were performed in 200 μL microcosms and incubated in a TECAN SPARK microtiter plate reader at 28°C using optical density measured at 600 nm (OD_600_).

### Comparative proteomics of *Flavobacterium* spp

Methods adapted from (*37, 74*) were combined. Briefly, 25 mL cell cultures (n=3) grown to an OD_600_ ∼0.6-1 were harvested by centrifugation at 3200 x g for 45 min at 4°C. Cells were resuspended in 20 mM Tris-HCl pH 7.8 and re-pelleted at 13000 x g for 5 min at 4°C. Cell lysis was achieved by boiling in 100 μl lithium dodecyl sulphate (LDS) buffer (Expedeon) prior to loading 20 μl onto a 4-20% Bis-Tris sodium dodecyl sulphate (SDS) precast gel (Expedeon). SDS-PAGE was performed with RunBlue SDS Running Buffer (TEO-Tricine) 1X (Expedeon) at 140 V for 5-10 min. Gels were stained with Instant Blue (Expedeon). A single gel band containing all the protein was excised. Gel sections were de-stained with 50 mM ammonium bicarbonate in 50% (v/v) ethanol, dehydrated with 100% ethanol, reduced and alkylated with Tris-2-carboxyethylphosphine (TCEP) and iodoacetamide (IAA), washed with 50 mM ammonium bicarbonate in 50% (v/v) ethanol and dehydrated with 100% ethanol prior to overnight digestion with trypsin. Samples were analysed by nanoLC-ESI-MS/MS using an Ultimate 3000 LC system (Dionex-LC Packings) coupled to an Orbitrap Fusion mass spectrometer (Thermo Scientific, USA) using a 60 min LC separation on a 25 cm column and settings as previously described (*75*). Resulting tandem mass spectrometry (MS/MS) files were searched against the relevant protein sequence database (*F. johnsoniae* UW101, UP000214645, *Flavobacterium* sp. OSR005 (Table SX) using MaxQuant with default settings and quantification was achieved using Label Free Quantification (LFQ). Statistical analysis and data visualisation of exoproteomes was carried out in Perseus (*76*).

### Bacterial genetics

To construct various XyGUL mutants, the method from (*77*) was adapted, as per our previous study (*36*). Briefly, fragments ∼1.5 kb in length upstream and downstream of the targeted genes were cloned into the plasmid pYT313 using the HiFi assembly kit (New England Biosciences). A full list of primers can be found in Table S1. Plasmid inserts were verified by Sanger sequencing. The resulting plasmids were transformed into the donor strain *E. coli* S17-1 λpir (S17-1 λpir) and mobilised into *F. johnsoniae* via conjugation: overnight (5 mL) *F. johnsoniae* wild type and pYT313-transformed S17-1 λ*pir* cultures were inoculated (20% v/v) into fresh CYE (5 mL) and incubated for a further 8 h. Cells were pelleted at 1800 x g for 10 min @ 22°C and washed in 1 mL CYE, and a 200 μL donor: recipient (CYE) suspension (1:1) was spotted onto CYE containing CaCl_2_ (10 mM) and incubated overnight at 28 °C. Biofilms were scraped from the agar surface and resuspended in 1 mL minimal A medium (no C source). Transconjugants were selected by spreading 5 to 100 μL aliquots on CYE containing erythromycin (100 μg mL^-1^). Colonies were restreaked onto CYE erythromycin and single homologous recombination events were confirmed by PCR prior to overnight growth in CYE followed by plating onto CYE containing 10% (w/v) sucrose to select for a second recombination event resulting in plasmid excision. To identify double homologous recombinants, colonies were replica plated onto CYE containing 10% (w/v) sucrose and CYE containing erythromycin. Erythromycin-sensitive colonies were screened by PCR.

For complementation of the *F. johnsoniae* Δ*xusCD* mutant, both genes and the 300-bp upstream region were cloned into pCP11 using the HiFi assembly kit. The insert was verified by Sanger sequencing and the plasmid was mobilised into DSM2064 via conjugation using S17-1 λ*pir* as the donor strain. The method was identical to that described above for transfer of the suicide plasmid, pYT313, except that 1 mL overnight cultures of donor and recipient were directly washed and resuspended in 200 μL CYE prior to spotting onto CYE containing CaCl_2_ (10 mM). Cells were scraped from the solid medium and transformants selected by creating a serial dilution (10^-1^ to 10^-5^) from the cell suspension and spotting 20 μL of each dilution onto CYE containing erythromycin (100 μg mL^-1^).

### Production and purification of recombinant GH5_4 homologs

Genes encoding the GH5_4 homologs (Fjoh_0774, BACOVA_02653, OSR005_04227 and OSR005_03871) lacking the N-terminal signal peptide and stop codon were amplified by PCR and ligated into the NdeI and XhoI sites of pET21a. Site-directed mutagenesis of the Trp252 residue in *Bo*GH5A was performed using the QuikChange II Site Directed Mutagenesis (SDM) Kit (Agilent Technologies) according to the manufacturer’s protocol.

For production of recombinant proteins, a single colony of *E. coli* BL21 (DE3) transformed with the desired plasmid was inoculated in 5 mL LB broth with 100 μg/mL ampicillin and shaken (220 rpm) at 37 °C overnight (16 h) before transfer to 1 L LB culture (in a 2 L conical flask) supplemented with 100 μg/mL ampicillin. Cultures were shaken at 37 °C at 220 rpm until an optical density at 600 nm (OD_600_) of ∼ 0.6 was reached. Following induction of gene expression with 0.4 mM (final concentration) IPTG, cells were incubated at 18°C overnight for a further 16 h before recovery by centrifugation at 8,000 x *g* for 15 min at 4°C. Pellets were resuspended in 30 mL binding buffer (25 mM HEPES pH 7.4, 1 M NaCl, 5 mM imidazole) and stored at -20 °C until purification. Cells were thawed and lysed by sonication. The lysate was centrifuged at 13,000 *x g* for 15 min at 4 °C and the supernatant was loaded onto a 5 ml chelating Sepharose column charged with nickel (II) sulphate pre-equilibrated with 50 mL of binding buffer. Following washes in binding buffer with increasing concentrations of imidazole, proteins were eluted with 25 mM HEPES pH 7.4, 400 mM Imidazole, 100 mM NaCl. Fractions containing the target protein (as identified by SDS-PAGE) were pooled and concentrated to a volume of 1-2 mL using a Vivaspin centrifuge concentrator (Sartorius) with a 30,000 kDa molecular weight cut off. The concentrated sample was loaded onto a size exclusion chromatography (SEC) (S200 16/60 Cytiva) column equilibrated in 50 mM Tris-HCl, 200 mM NaCl, 10 % (w/v) glycerol and protein was separated at a flow rate of 0.5 mL min^-1^. Purity of peak fractions was analysed by SDS-PAGE and protein was stored at -20 °C until required.

### Enzymatic assays of recombinant glycoside hydrolases

Purified recombinant GH5_4 homologs were screened for enzyme activity using the 3,5-Dinitrosalicylic Acid Assay (DNSA) method (*78*). Briefly, for enzyme kinetics between 10-250 nM purified recombinant protein (n=3) was incubated with decreasing concentrations (starting from 8 mg mL^-1^) of XyG. At each time point a subsample was taken and mixed with a stop solution (DNSA working reagent containing 10 mg ml^-1^ glucose), prior to boiling at 95°C for 15 min to develop the colour. To calculate the initial maximum velocity of the reaction (*V*_o_), at least five measurements were taken within the linear kinetics range. Absorbance at 575 nm was quantified. A standard curve (n=3) against known concentrations of glucose was used to convert A575 to the amount of freely available reducing ends produced during cleavage of the beta-glucan backbone of XyG. All assays were typically repeated with two separate batches of protein. For screening the promiscuous activity of OSR005-1, OSR005-2, *Bo*GH5A, *Bo*W252A or *Bo*W252G, 1 μM of protein was incubated with 4 mg mL^-1^ polysaccharide for 30 min.

### Comparative genomics and metagenomics

The online platform IMG/JGI (*79*) was used to conduct most comparative genomics analyses described in this study . Genomes and metagenomes were stored in genome sets (detailed in Tables S2 and S3), and BLASTP searches (E-value e^-40^) were performed using the “jobs function” using either Fjoh_0774 or a homologue (IMG gene id; 2644426200) of the commercial recombinant endoxyloglucanase (GH5_4) from *Paenibacillus* sp. (Megazyme, CAS - 76901-10-5). The latter was used as it represents a sequence from outside GH5_4 subclade 2. For the metagenome searches, retrieved open reading frames (ORFs, n=7136) were locally aligned (BLASTP) against all GH5 ORFs (n=1246) deposited in the CAZy database (CaZydb) (*80*). The estimated gene copy index (calculated by using the average read coverage depth across a given contig) provided by IMG/JGI was used to calculate the relative abundance for each retrieved ORF. To aid with the identification and organisation of PUL, the PULDB online platform was also utilised (*81*).

## Author contributions

IL conceived and designed the project with consultation from AH. HM, LR, LMK, PF, AAB, AQ, AH and IL carried out the experimental work. AM performed the comparative proteomics analysis. IL performed the comparative genomics and metagenomics. HM, LR, AH and IL wrote the paper with TD, DN and SA providing feedback.

## Acknowledgements

We thank the Warwick Proteomics Research Facility, namely Dr Cleidiane Zampronio and Dr Andrew Bottrill, for their assistance in generating and processing the mass-spectrometry data. This study was funded by a Biotechnology and Biological Sciences Research Council (BBSRC) Discovery Fellowship (award BB/T009152/1) and a Royal Society University Research Fellowship (award URF\R1\221708) to IL. AH also acknowledges the support of a Royal Society University Research Fellowship (award URF\R1\191548). HM and PF were supported by The University of Sheffield through a Faculty of Science funded PhD studentship and Summer Undergraduate Research Experience (SURE) scheme, respectively. AQ was supported by a summer studentship funded by a Rank Prize Fund New Lecturer Award to IL.

