## Supplementary figures for "Hybrid xyloglucan utilisation loci are prevalent among plant-associated Bacteroidota"

### Slide 1
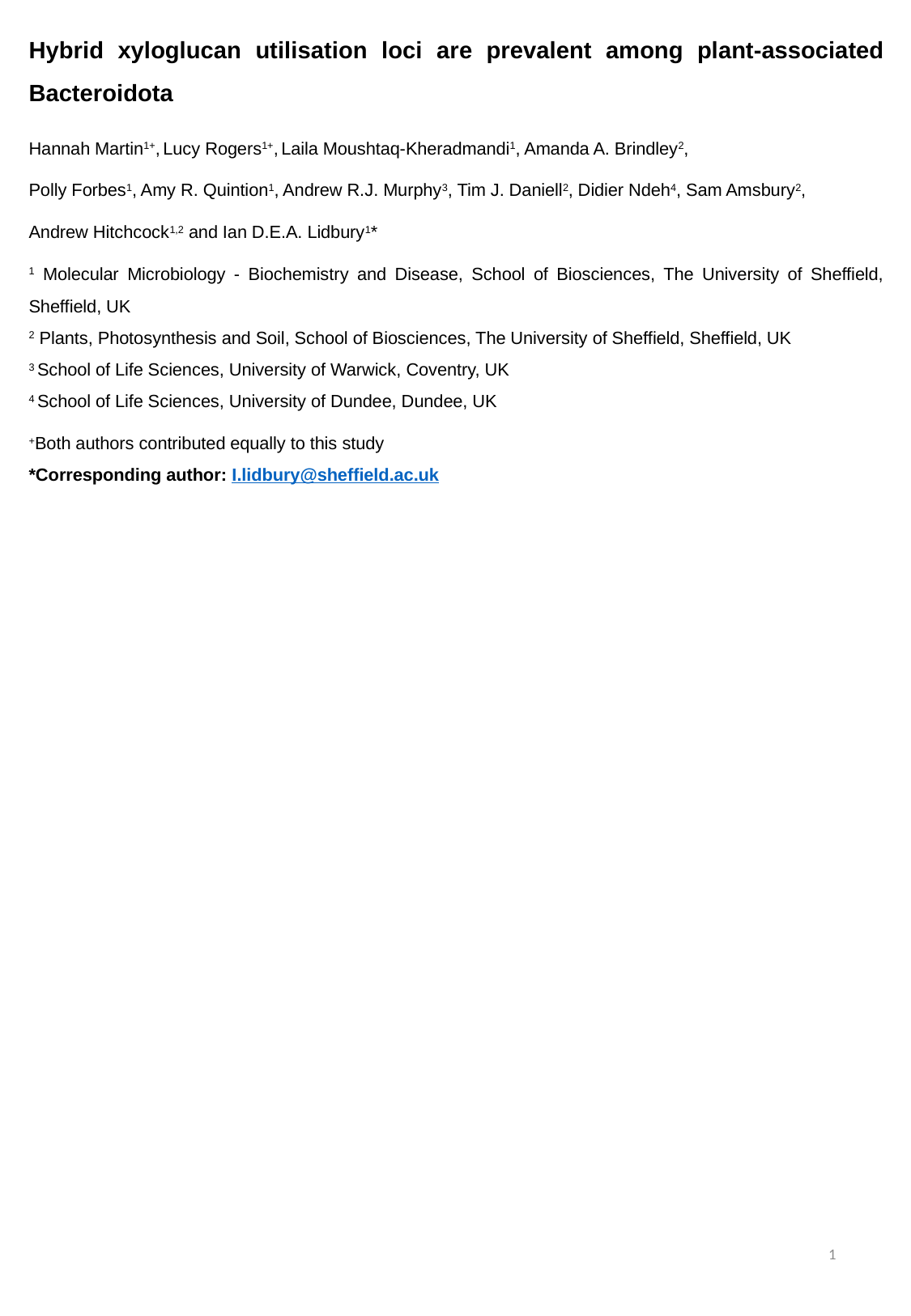

Hybrid xyloglucan utilisation loci are prevalent among plant-associated Bacteroidota
Hannah Martin^1+^, Lucy Rogers^1+^, Laila Moushtaq-Kheradmandi^1^, Amanda A. Brindley^2^,
Polly Forbes^1^, Amy R. Quintion^1^, Andrew R.J. Murphy^3^, Tim J. Daniell^2^, Didier Ndeh^4^, Sam Amsbury^2^,
Andrew Hitchcock^1,2^ and Ian D.E.A. Lidbury^1^*
^1^ Molecular Microbiology - Biochemistry and Disease, School of Biosciences, The University of Sheffield, Sheffield, UK
^2^ Plants, Photosynthesis and Soil, School of Biosciences, The University of Sheffield, Sheffield, UK
^3^ School of Life Sciences, University of Warwick, Coventry, UK
^4^ School of Life Sciences, University of Dundee, Dundee, UK
^+^Both authors contributed equally to this study
1

### Slide 2
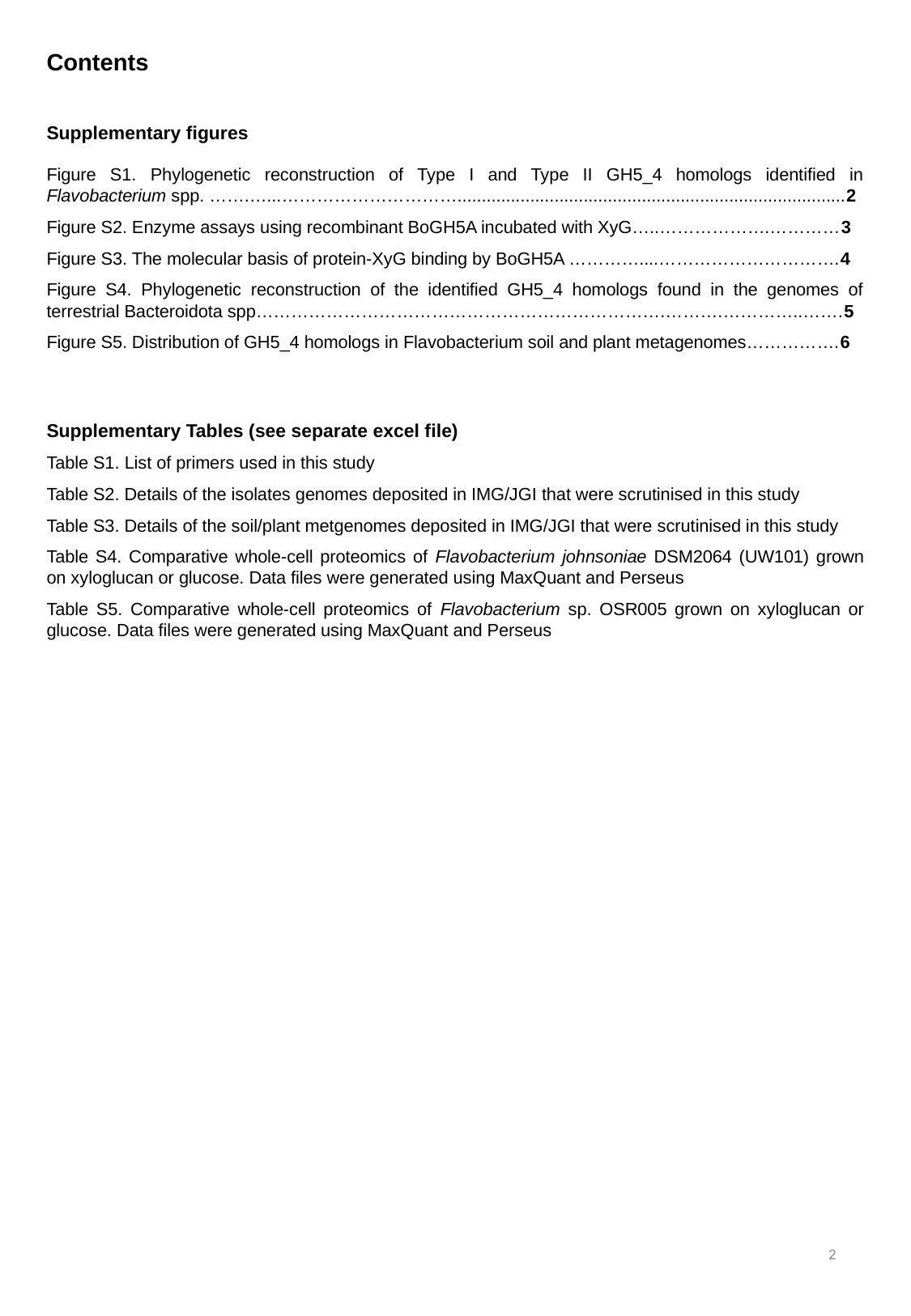

Contents
#
Supplementary figures
Figure S1. Phylogenetic reconstruction of Type I and Type II GH5_4 homologs identified in Flavobacterium spp. …….…...…………………………................................................................................2
Figure S2. Enzyme assays using recombinant BoGH5A incubated with XyG…..……………….…………3
Figure S3. The molecular basis of protein-XyG binding by BoGH5A …………....………………………….4
Figure S4. Phylogenetic reconstruction of the identified GH5_4 homologs found in the genomes of terrestrial Bacteroidota spp…………………………………………………………….……….…………..…….5
Figure S5. Distribution of GH5_4 homologs in Flavobacterium soil and plant metagenomes…………….6
Supplementary Tables (see separate excel file)
Table S1. List of primers used in this study
Table S2. Details of the isolates genomes deposited in IMG/JGI that were scrutinised in this study
Table S3. Details of the soil/plant metgenomes deposited in IMG/JGI that were scrutinised in this study
Table S4. Comparative whole-cell proteomics of Flavobacterium johnsoniae DSM2064 (UW101) grown on xyloglucan or glucose. Data files were generated using MaxQuant and Perseus
Table S5. Comparative whole-cell proteomics of Flavobacterium sp. OSR005 grown on xyloglucan or glucose. Data files were generated using MaxQuant and Perseus
2

### Slide 3
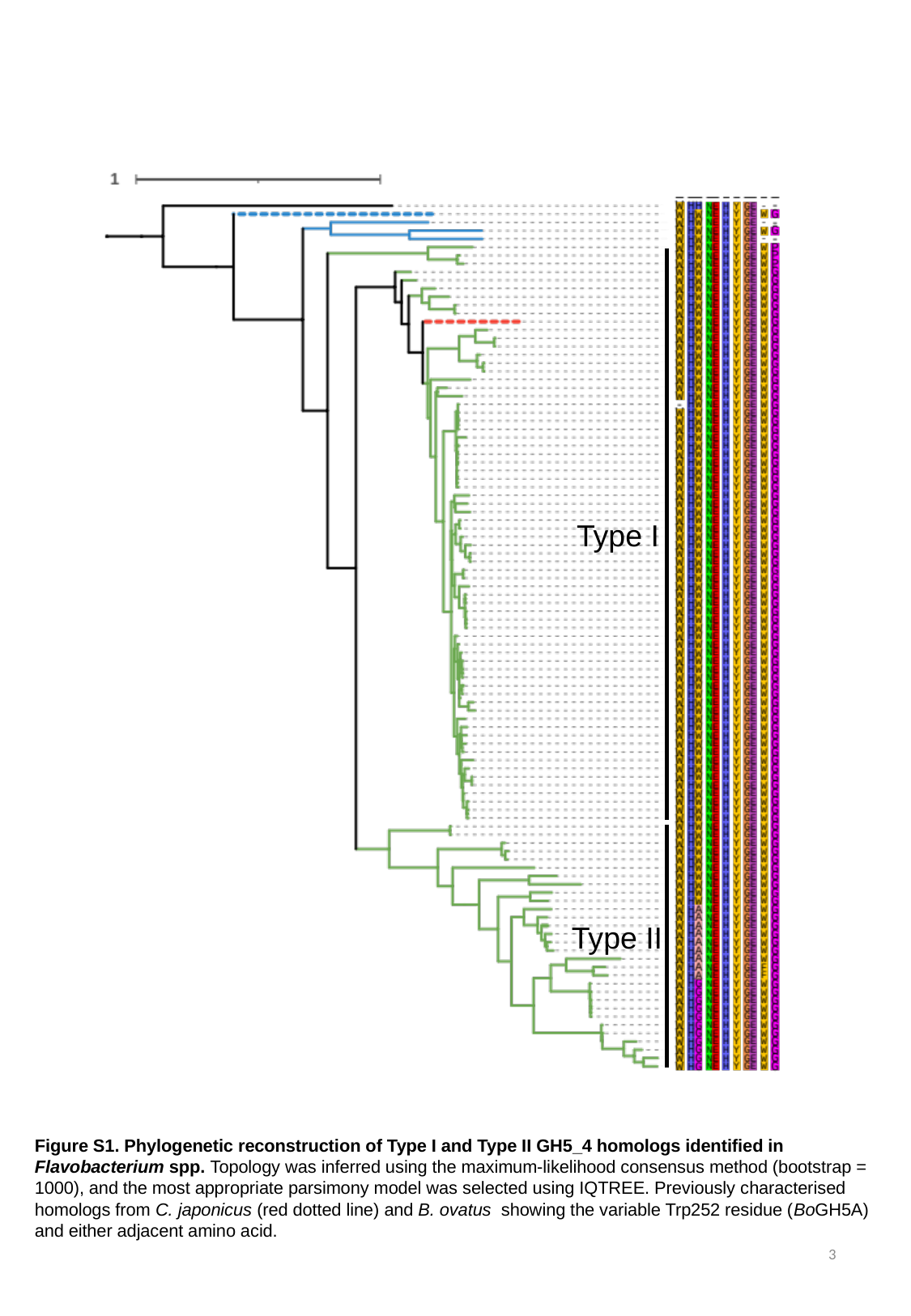

Type I
Type II
Figure S1. Phylogenetic reconstruction of Type I and Type II GH5_4 homologs identified in Flavobacterium spp. Topology was inferred using the maximum-likelihood consensus method (bootstrap = 1000), and the most appropriate parsimony model was selected using IQTREE. Previously characterised homologs from C. japonicus (red dotted line) and B. ovatus showing the variable Trp252 residue (BoGH5A) and either adjacent amino acid.
3

### Slide 4
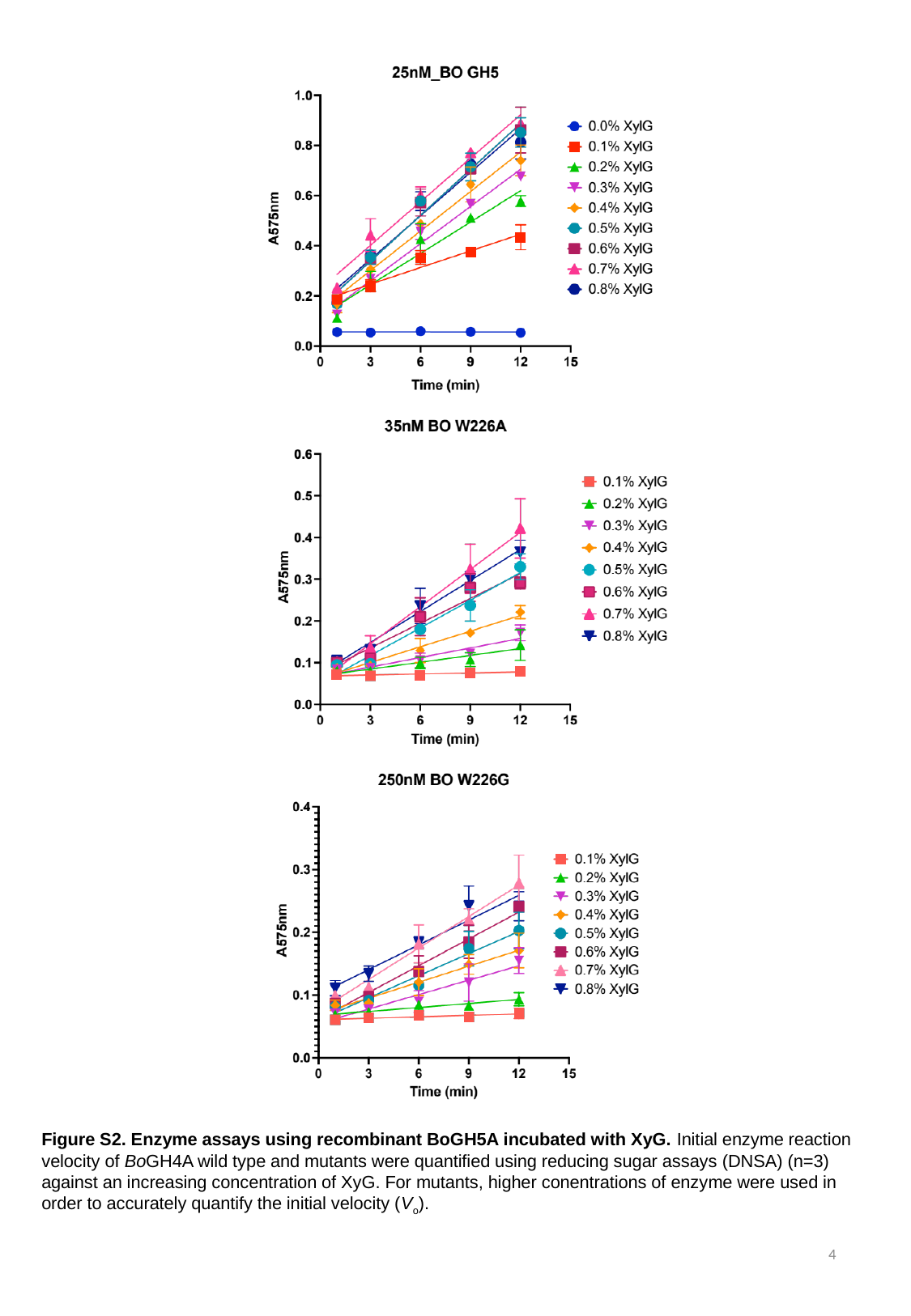

Figure S2. Enzyme assays using recombinant BoGH5A incubated with XyG. Initial enzyme reaction velocity of BoGH4A wild type and mutants were quantified using reducing sugar assays (DNSA) (n=3) against an increasing concentration of XyG. For mutants, higher conentrations of enzyme were used in order to accurately quantify the initial velocity (V_o_).
4

### Slide 5
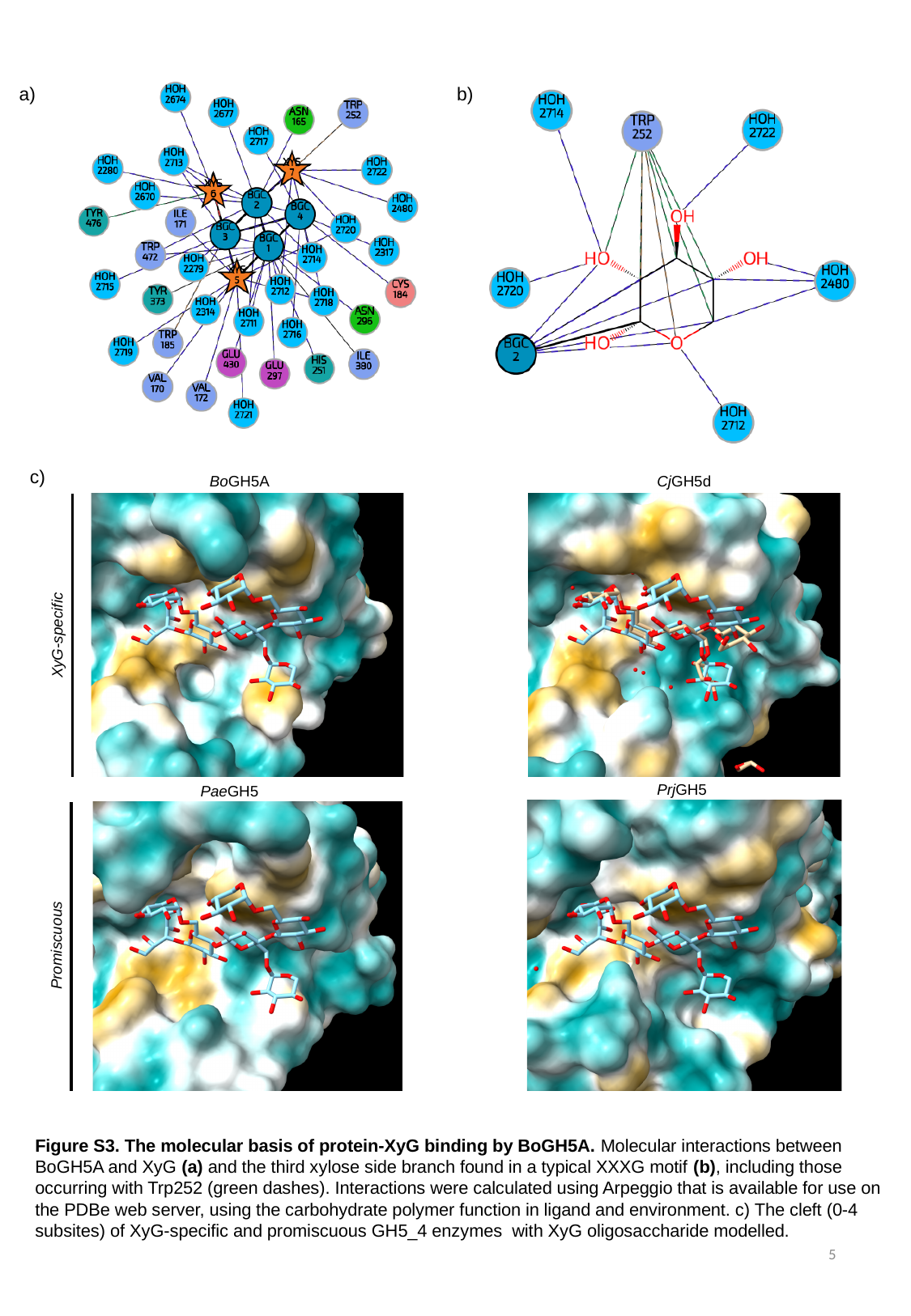

a)
b)
b)
c)
BoGH5A
CjGH5d
XyG-specific
PrjGH5
PaeGH5
Promiscuous
Figure S3. The molecular basis of protein-XyG binding by BoGH5A. Molecular interactions between BoGH5A and XyG (a) and the third xylose side branch found in a typical XXXG motif (b), including those occurring with Trp252 (green dashes). Interactions were calculated using Arpeggio that is available for use on the PDBe web server, using the carbohydrate polymer function in ligand and environment. c) The cleft (0-4 subsites) of XyG-specific and promiscuous GH5_4 enzymes with XyG oligosaccharide modelled.
5

### Slide 6
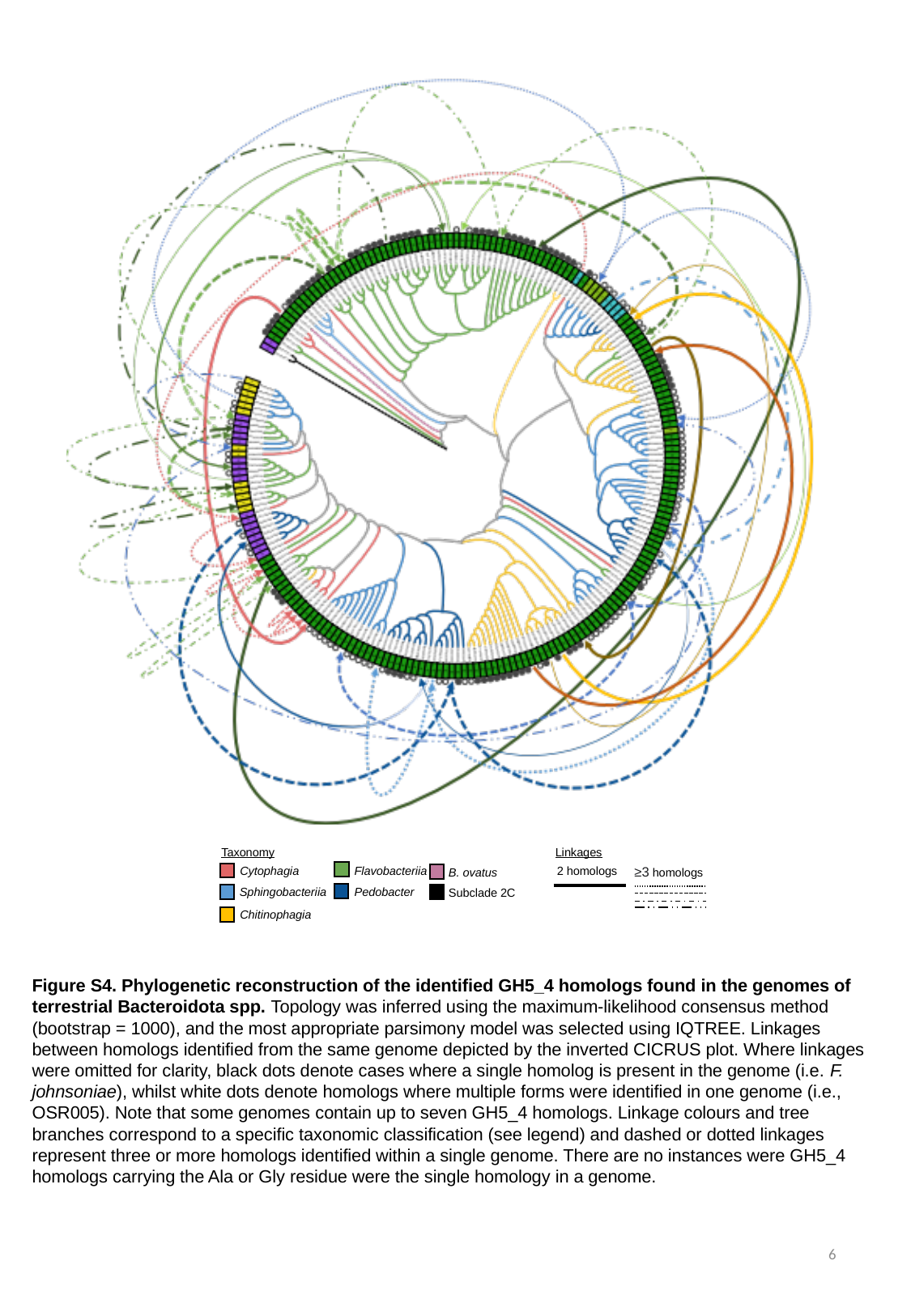

Taxonomy
Linkages
Cytophagia
2 homologs
≥3 homologs
Flavobacteriia
B. ovatus
Sphingobacteriia
Pedobacter
Subclade 2C
Chitinophagia
Figure S4. Phylogenetic reconstruction of the identified GH5_4 homologs found in the genomes of terrestrial Bacteroidota spp. Topology was inferred using the maximum-likelihood consensus method (bootstrap = 1000), and the most appropriate parsimony model was selected using IQTREE. Linkages between homologs identified from the same genome depicted by the inverted CICRUS plot. Where linkages were omitted for clarity, black dots denote cases where a single homolog is present in the genome (i.e. F. johnsoniae), whilst white dots denote homologs where multiple forms were identified in one genome (i.e., OSR005). Note that some genomes contain up to seven GH5_4 homologs. Linkage colours and tree branches correspond to a specific taxonomic classification (see legend) and dashed or dotted linkages represent three or more homologs identified within a single genome. There are no instances were GH5_4 homologs carrying the Ala or Gly residue were the single homology in a genome.
6

### Slide 7
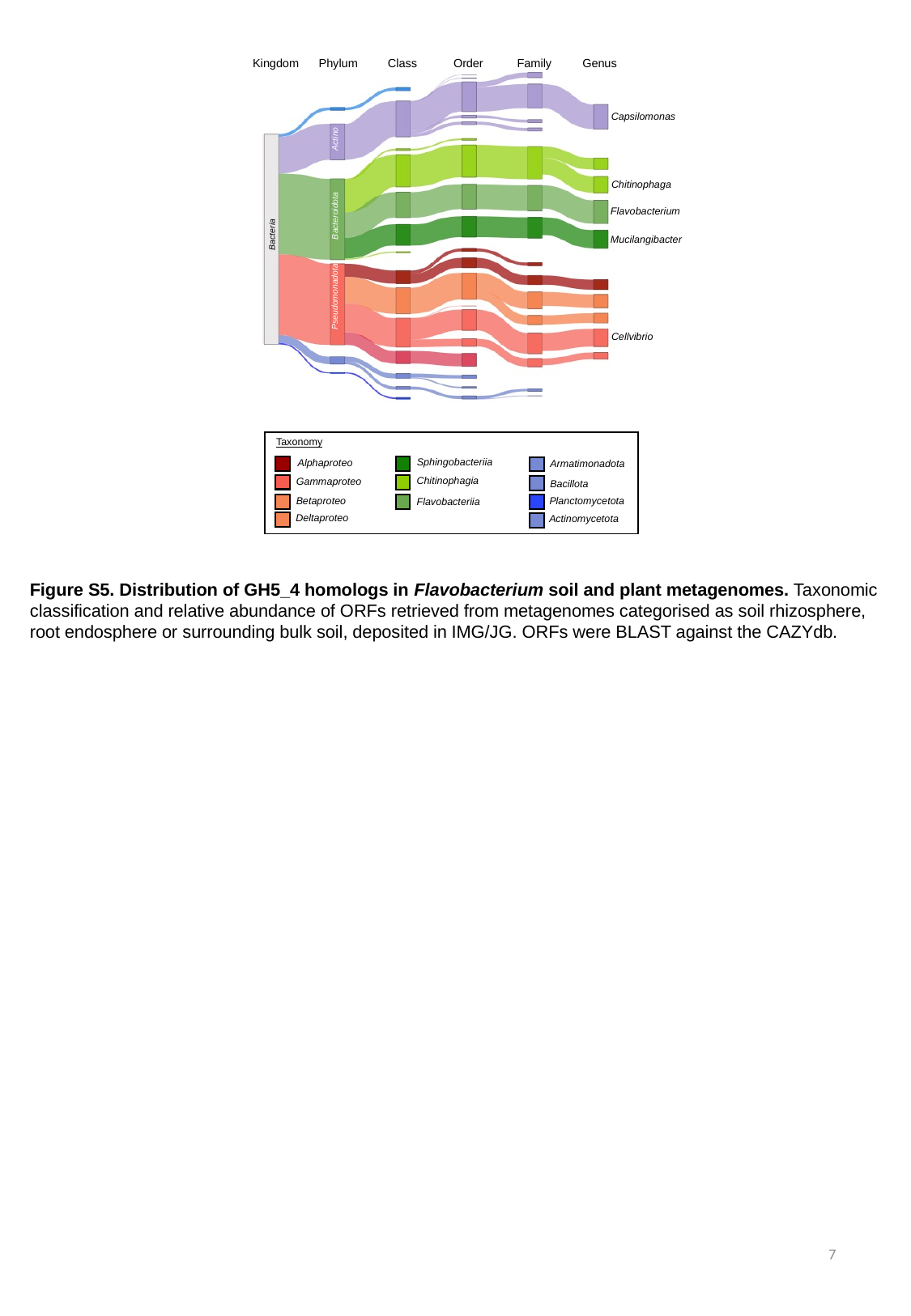

Kingdom
Phylum
Order
Family
Genus
Class
Figure S3
Capsilomonas
Actino
Chitinophaga
Flavobacterium
Bacteroidota
Bacteria
Mucilangibacter
Pseudomonadota
Cellvibrio
Taxonomy
Sphingobacteriia
Alphaproteo
Armatimonadota
Chitinophagia
Gammaproteo
Bacillota
Betaproteo
Planctomycetota
Flavobacteriia
Deltaproteo
Actinomycetota
Figure S5. Distribution of GH5_4 homologs in Flavobacterium soil and plant metagenomes. Taxonomic classification and relative abundance of ORFs retrieved from metagenomes categorised as soil rhizosphere, root endosphere or surrounding bulk soil, deposited in IMG/JG. ORFs were BLAST against the CAZYdb.
7
